# Amphibian diversity and abundance in ponds is lower in exotic plantations than native forests

**DOI:** 10.1101/074302

**Authors:** Maider Iglesias-Carrasco, Iraide Artexte, Carlos Cabido, Aitor Larrañaga

## Abstract

What effect do tree plantations have on the diversity of native organisms? Some studies show that plantations reduce the diversity and abundance of certain taxa, while other studies suggested that plantations help to conserve biodiversity. Pine and eucalyptus plantations are among the most widespread exotic plantations worldwide, and they have negative effects on many taxa. But how do they affect amphibian diversity and abundance? We barely know. We therefore tallied up the number of amphibian taxa and their abundance from 18 ponds in patches of native oak forests, pine or eucalypt plantations. We also quantified water quality by measuring its physicochemistry and identifying the macroinvertebrates present in each pond. There were significantly fewer amphibian species in tree plantations than in native forest. Compared to native forest, the total density of amphibians was also significantly lower in eucalypt, but not pine, plantations. Species varied in the effects of plantations on their presence and abundance. We suggest that the decline in the presence and abundance of amphibians in plantations is linked to the physicochemical of pond water, combined with the relatively low presence of invertebrate. It seems likely that earlier desiccation, greater toxicity, and poor quality detritus in ponds in plantation are key drivers of species decline. The effects of these drivers are expected to worsen as climate change continues.

## Introduction

All over the world vast areas of native forest have been converted into exotic tree plantations. In total, 7% of the global forested area is occupied by plantations [1], which are mainly used to produce paper, timber or charcoal [2]. Plantations have a higher density, but lower diversity, of trees than natural forests [3]. Many studies have shown that plantations reduce the number of species of important animal groups, such as arthropods and birds [4,5]. But in extreme cases where natural forests become scarce, plantations can paradoxically contribute to the conservation of diversity [6–8]. Eucalypt (mostly *Eucalyptus globulus* Labill.) and pine plantations (*Pinus sp.*) are the two most common types of exotic plantations worldwide. We now have information about how pines and eucalypts affect many taxa, including birds [9], mites [10], fish [11], lizards [12] and even native vegetation [13]. In contrast, how plantations affect amphibians is practically unknown (but see [14,15]). Here we address this gap in knowledge.

As with many non-native plants, eucalypt plantations release toxic substances into the soil [16] and waterways [17], reduce water yields and change soil characteristics [18]. Widely reported declines in stream assemblages, such as stream macroinvertebrates and fungi, in waterways in the catchments of eucalypt and pine plantations [19–21] have been attributed to the toxicity and low quality of their leaf litter. We expect these effects to be stronger in stationary (lentic) water systems, as water quality decline is exacerbated due to the steady accumulation of leaf litter and the lack of water renewal [22].

Globally, amphibians are among the most threatened animal groups. This is largely due to habitat loss [23], which is particularly devastating as amphibians often have low mobility and high philopatry [24]. Their highly permeable skin also makes them sensitive to toxic substances [25]. Consequently, small changes in the chemical characteristics of their terrestrial or aquatic habitats (depending on the life history phase) can have major effects on amphibian survival and breeding success. For example, the introduction of some exotic plants alters native forest amphibian communities due to seemingly small modifications, such as a changes in the temperature in their preferred microhabitats [26]. Small, ephemeral ponds are common in forests and usually support a high diversity and abundance of amphibians. Many amphibians breed in such ponds, stay there until they complete larval development, and then emerge to forage and hibernation/estivation in nearby terrestrial habitat [27]. Lower species diversity and abundance in plantations could be due to negative effects of leachates from their leaf litter on water quality and therefore larval survival and/or adult breeding success; or to plantations lowering the survival of juveniles and adults while on land.

We investigated how *Eucalypt* and *Pine* plantations affect the diversity and relative density of amphibian adults *and* larvae. Most European amphibians have terrestrial adults that move to nearby ponds to breed. We therefore focused our sampling efforts in the vicinity of, and in, ponds. We first characterized the amphibian assemblages in ponds under native forests and *Eucalypt* and *Pine* plantations. We then measured physical, chemical and biological properties of these ponds to determine which environmental characteristics might affect amphibian diversity and abundance. We tested whether:

1. replacement of native forest by pine and eucalypt exotic plantations reduces the diversity and density of amphibian species;
2. the assemblage of macroinvertebrates in ponds predicts amphibian diversity and density, because these assemblages are related to long-term water quality and are themselves a food resource for amphibians;
3. water chemistry, wetland vegetation and the size/depth of ponds predict amphibian diversity and density.

## Material and methods

### Study site

We collected data at 18 ponds in Atlantic watersheds of the Basque Country: six under native deciduous forest patches (*Quercus robur* L.), six under eucalypt plantations *(Eucalyptus globulus)* and another six under pine plantations *(Pinus radiata D.Don)*. The ponds were totally surrounded for at least 400 meters by the corresponding habitat type. The climate of the study area is mesotemperate oceanic, with an average annual temperature of 11.6–13.1ºC and an average annual precipitation of 1200 to >2000 mm [28].

### Sampling of amphibians and macroinvertebrates

We sampled each pond twice, in mid-March and in late May 2015, to increase the likelihood of finding both winter and spring/summer breeding amphibians. Each sampling bout was less than a week to reduce confounding effects of weather or time in the life cycle on habitat differences. All ponds were dipnetted by MI-C for invertebrates and amphibians (larvae and adults) in a standardized way (effort: 1 minute m^−2^; net size: 1 mm mesh, 26 × 21 cm frame). We identified all amphibian species *in situ*, recorded the number of individuals and determined their sex and life cycle stages (larvae, metamorphic, juvenile or adult). We then released them back into the pond. We stored macroinvertebrates in 70% ethanol and transported them to the laboratory for identification to the family level following Tachet et al., 2010. For our statistical analyses we used two biological index based on the sensitivity of invertebrate families to water contamination: the IBMWP (Alba-Tercedor et al., 2002) and the Iberian Average Score Per Taxon (IASPT) [31], calculated as the division of the IBMWP and the number of different taxa.

### Environmental characteristics

At each pond we measured 8 variables that seem to be important drivers for amphibian and macroinvertebrate diversity in aquatic habitats [32–36]. We measured the water temperature, pH, and conductivity with field WTW multi-parametric sensors. To measure light penetration in water, we used a LI-COR Li-250A light meter placed at the center of the pond and then expressed the pond’s turbidity as the light extinction coefficient (K). We also estimated the pond’s area and average depth (mean of five measures along the length of the pond). For this, using the longest axis of the pond as reference (*a*), we defined a number of equidistant and perpendicular transects (*b*_*1*_ to *b*_*n*_) (total transects depended on the irregularity of the pond: range 2–8). The area of the pond was estimated as *a*\*average (*b*_*1*_ to *b*_*n*_). Depth was recorded every 15 cm along those transects and we used the mean using all the depths computed. Finally, we estimated the percentage of the pond covered by submerged and emergent aquatic vegetation.

### Statistical analyses

We used linear models to test for differences in the measured variables among the three habitat types. When necessary, data were log-transformed to meet model requirements. Multiple regressions by forward selections were performed to predict amphibian richness and total amphibian density [37]. For that, we performed linear models between the dependent variable and each independent variable alone and we retained those independent variables that fit best (lowest AIC). We then added new independent variables that reduced the most the previous AIC values. We stopped adding new variables when reductions of the AIC were smaller than a value of 2 [37].

## Results

### Biodiversity and abundance of amphibians

We found 5494 individuals (larvae and adults) from 7 species in ponds under native forests, 885 individuals of 2 species under pine plantations and 168 individuals of 4 species under eucalypt plantations (Fig 1a). Only *Lissotriton helveticus* (Razoumowsky, 1789) was detected in all the sampled ponds (Table 1). By contrast, *Alytes obstetricans* (Laurenti, 1768)*, Rana dalmatina* Fitzinger, 1838 and *Triturus marmoratus* (Latreille, 1800), appeared only in native forests, while *Rana temporaria* Linnaeus, 1758, *Pelophylax perezi* (López Seoane, 1885) and *Salamandra salamandra* (Linnaeus, 1758) appeared in both native forests and eucalypt plantations (Table 1). We found both adults and larvae of *L. helveticus*; only larvae of *A. obstetricans*, *R. dalmatina, Rana temporaria*, *S. salamandra*; and only adults of *P. perezi* and *T. marmoratus* (Table 1). The number of species differed among the three habitat types (F2,15 = 9.95, p = 0.002, Fig. 1a). There were significantly more species in ponds in native forests (median: 4) than in either pine (1.5) or eucalypt plantations (2) (Fig 1a), but no difference between the two plantation types (Tukey HSD test: p = 0.70). There were also differences in total density between native forests and plantations (F_2,15_ = 4.29, p= 0.034, Fig 1b). Amphibian density was highest in ponds in native forests (mean ± SE: 72.53 ± 32.9 no. m^−2^), intermediate in pine plantations (17.60 ± 7.52 no.m^−2^) and lowest in eucalypt plantations (8.15 ± 5.53 no.m^−2^), but the only significantly pairwise different was between native forests and eucalypt plantations (Tukey HSD: p = 0.026). Looking at each species separately, only the densities of *A. obstetricans* and *R. temporaria* varied significantly with habitat type. *A. obstetricans* only inhabited native forests (1.77 ± 0.91 no.m^−2^) (Table 1). *R. temporaria* occurred in all three habitats but at significantly different densities. The density was highest in oak forest (62.99 ± 33.26 no.m^−2^), intermediate in pine plantations (10.37 ± 6.34 no.m^−2^) and lowest in eucalypt plantations (0.11 ± 0.11 no.m^−2^), although the only significant pairwise difference was between oak forests and eucalypt plantations (Tukey HSD: p = 0.014).

**Figure 1:**
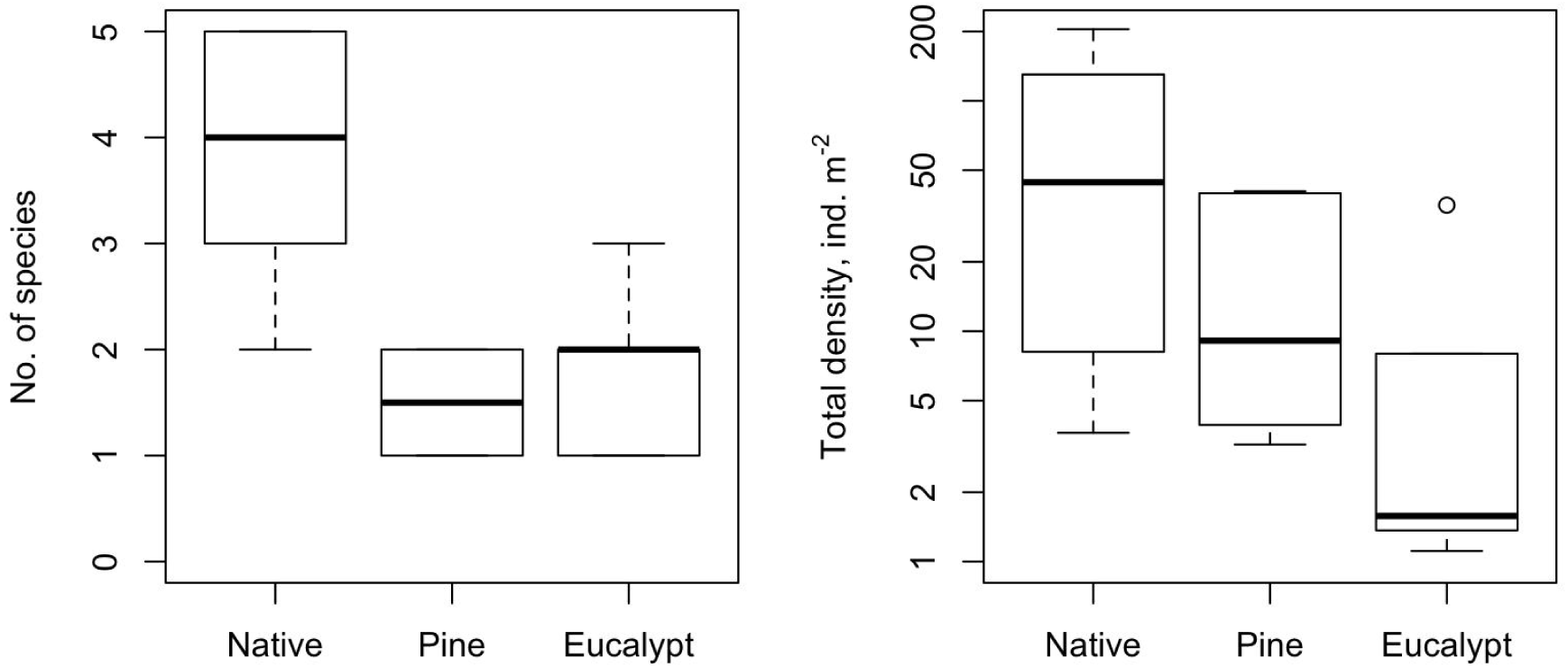
(a) Number of species and (b) total density of individuals per pond. ANOVA results for Richness: F_2,15_ = 9.95, p = 0.002; Tukey HSD comparison: Native > Eucalypt = Pine, and for Total density: F_2,15_ = 4.29, p = 0.034; Tukey HSD: Native > Eucalypus (the other pairwise tests are non-significant). Note the logarithmic y-axis for total density.

**Table 1:**
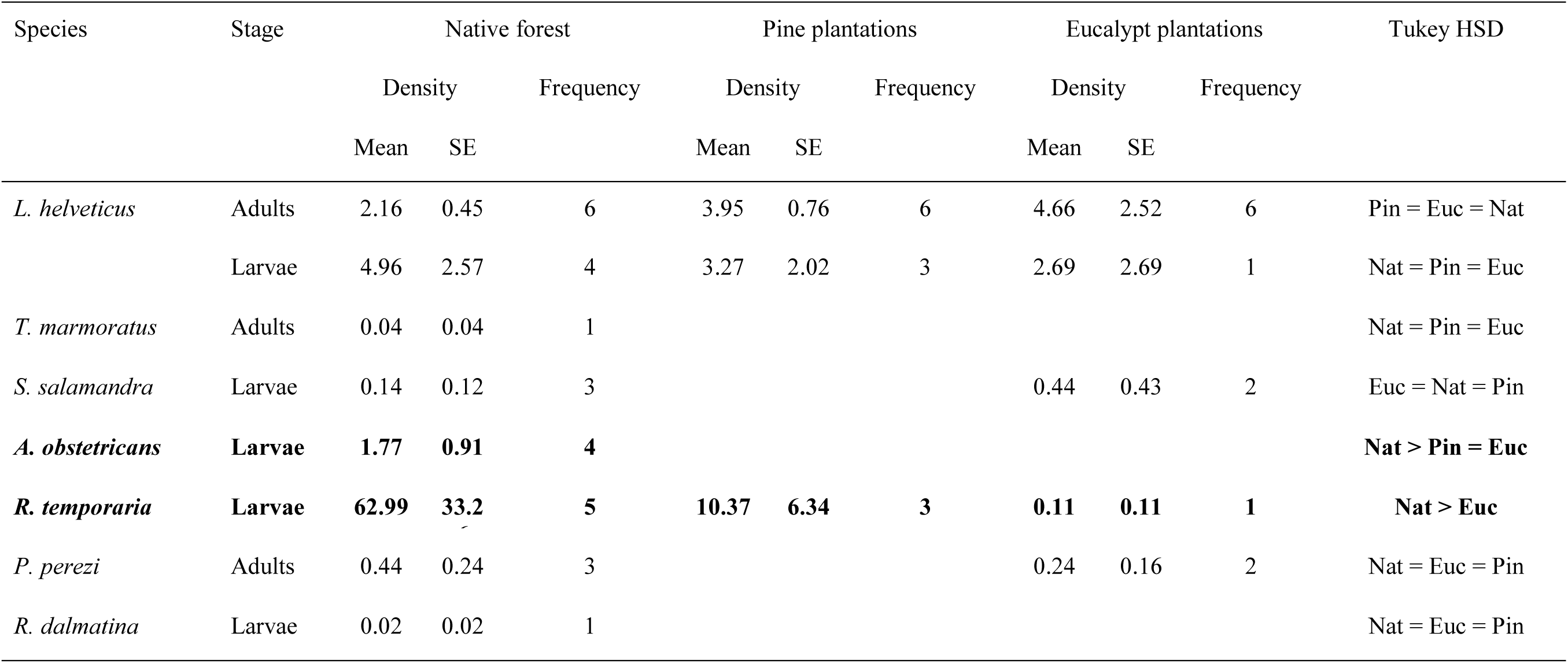
Density (mean ± SE, no. m^−2^) and frequency (out of 6 ponds) for each amphibian species separated in adults and larvae. Comparisons among the three habitat types after Tukey HSD tests are shown. In bold are species that differ in density among habitats. (N = 6 ponds per habitat type).

### Environmental characteristics of ponds and relationship with amphibians

Ponds in native forests were significantly deeper (23.55 ± 6.85 cm) than those in pine (7.79 ± 0.83 cm) and eucalypt plantations (8.44 ± 2.22 cm) (Table 2). Similarly, pond surface area was bigger in native forests (22 ± 5.90 m^2^) than in pine (7.16 ± 0.20 m^2^) and eucalypt (5.95 ± 1.59 m^2^) plantations (Table 2). Although submerged, emerged and terrestrial vegetation cover varied widely among habitats (range of average values: 0.8–23.3, 1.7–10.0 and 29.2–75.0, respectively), there were no significant differences (Table 2). Variation in water physicochemistry (pH, temperature, conductivity) was smaller and it did not differ significantly among habitat types (Table 2). For macroinvertebrates, only the IBMWP, the index of sensitivity of invertebrate families to water contamination, was significant different about habitat types: highest in native forests (56.67 ± 9.85), intermediate in pine plantations (31.83 ± 6.26) and lowest in eucalypt plantations (23.33 ± 6.77) (Tukey HSD: p = 0.022) (Table 2).

**Table 2:** Mean and SE of descriptors of the ponds sampled. Comparisons after Tukey HSD tests are shown for each variable. In bold are the variables with significant differences among habitat types.

| Descriptor | Native forest |  | Pine plantations |  | Eucalypt plantations |  | Tukey HSD |
| --- | --- | --- | --- | --- | --- | --- | --- |
|  | Mean | SE | Mean | SE | Mean | SE |  |
| **Mean water depth (cm)** | **23.55** | **6.85** | **7.79** | **0.83** | **8.44** | **2.22** | **Nat > Euc = Pin** |
| **Area (m<sup>2</sup>)** | **22.69** | **5.90** | **7.16** | **0.20** | **5.95** | **1.59** | **Nat > Pin = Euc** |
| Submerged vegetation (%) | 23.33 | 11.38 | 0.83 | 0.83 | 20.00 | 10.88 | Nat = Euc = Pin |
| Emerged vegetation (%) | 1.67 | 1.05 | 1.67 | 1.05 | 10.00 | 4.28 | Euc = Nat = Pin |
| Terrestrial vegetation (%) | 73.33 | 14.70 | 29.17 | 12.81 | 75.00 | 11.55 | Euc = Nat = Pin |
| pH | 7.34 | 0.15 | 6.74 | 0.45 | 7.35 | 0.26 | Euc = Nat = Pin |
| Temperature (°C) | 11.67 | 2.61 | 8.18 | 1.28 | 10.18 | 0.55 | Nat = Euc = Pin |
| Conductivity (µS cm<sup>-1</sup>) | 136.52 | 41.15 | 91.53 | 21.25 | 147.17 | 42.43 | Euc = Nat = Pin |
| Turbidity (K) | 6.40 | 1.01 | 6.08 | 1.23 | 8.69 | 1.33 | Euc = Nat = Pin |
| Invertebrate taxa (no. families) | 12.00 | 1.86 | 7.33 | 1.12 | 4.83 | 0.87 | Nat = Pin = Euc |
| Invertebrate predator taxa (%) | 48.88 | 3.75 | 59.72 | 10.19 | 52.50 | 12.44 | Pin = Euc = Nat |
| **IBMWP** | **56.67** | **9.85** | **31.83** | **6.26** | **23.33** | **6.77** | **Nat > Euc** |
| IASPT | 4.51 | 0.36 | 4.18 | 0.29 | 4.56 | 0.59 | Nat = Euc = Pin |

Finally, the best model to explain the species richness of amphibians mainly included habitat type (93.0% of the variance explained; p < 0.001), but also conductivity (4.5%; p < 0.001) and submerged vegetation cover (0.7%; p = 0.0499), with less than 2% left unexplained. In contrast, amphibian abundance was explained by habitat type (36.4% of the variance explained; p = 0.002), size of the pond (23.4%, p = 0.003), terrestrial vegetation cover (11.6%, p = 0.024) and by the emerged vegetation cover (7.7%; p = 0.058; it was retained by the AIC criterion, despite not being significant in the ANOVA) (Table 3), with 21% left unexplained.

**Table 3:** Model supported by AIC for Mean density and Total richness of amphibians in the ponds.

| Response variable | Variable | Transformation | D | SS | MS | F | P | Variance | AIC | Comparison/Slope |
| --- | --- | --- | --- | --- | --- | --- | --- | --- | --- | --- |
| Total amphibian richness | Land use |  | 2 | 30.5 | 1.53 | 32.13 | <0.001 | 93.0 | 22.05 | Nat > Euc = Pin |
| | Conductivity ($\mu\text{S cm}^{-1}$) | LN | 1 | 1.46 | 1.46 | 30.76 | <0.001 | 4.5 | 5.91 | - |
|  | Submerged vegetation (%) | LN | 1 | 0.22 | 0.22 | 4.67 | 0.0499 | 0.7 | 2.38 | + |
|  | Residuals |  | 1 | 0.62 | 0.05 |  |  | 1.9 |  |  |
| Mean amphibian density | Land use |  | 2 | 16.28 | 8.14 | 10.41 | 0.002 | 36.4 | 67.33 | Nat > Euc |
| | Size ($\text{m}^2$) | LN | 1 | 10.45 | 10.45 | 13.36 | 0.003 | 23.4 | 61.09 | - |
|  | Terrestrial vegetation (%) | LN | 1 | 5.19 | 5.19 | 6.63 | 0.024 | 11.6 | 56.98 | + |
|  | Emerged vegetation (%)* | Non-transf. | 1 | 3.44 | 3.44 | 4.39 | 0.058 | 7.7 | 53.36 | - |
|  | Residuals |  | 1 | 9.39 | 0.78 |  |  | 21.0 |  |  |

\*Although non-significant in the ANOVA, AIC supported the inclusion of this variable in the model.

## Discussion

Species richness of amphibians was lower in ponds in both pine and eucalypt plantations than in native forests, while amphibian density was also significantly lower in eucalypt, but not pine, plantations than in native forests. Far more of the variance among ponds in species richness and density was explained by the habitat type (93 and 36%, respectively) than the specific properties of the ponds that we measured (e.g. pond size, depth and aspects of water quality). This suggests that variables that we did not measure that are themselves affected by the habitat type are responsible for the observed differences in amphibian diversity (e.g. toxin levels). We conclude that the replacement of natural forests by plantations is harmful for amphibians, albeit less so when the plantation is pine rather than eucalyptus. Our results also suggest that even the small number of forest patches sampled, the effect of the type of habitat is strong enough to be detected. The heterogeneity of our samplings was controlled, in part, by choosing small and temporary ponds in all the habitats. So that, small sample sizes combined with a low heterogeneity among ponds, seem to be enough to determine habitat modification effects on amphibian diversity.

### Plantation effects on water and resource quality

We characterized the physicochemistry of the ponds to look at how plantations affect water quality, but we did not detect any differences among the three habitat types. At first glance these findings are unexpected as eucalypt leachates can cause a marked decrease in oxygen level and pH, and an increase in conductivity in only a few days [22]. However, leaves from a range of tree species from alder, to eucalypt and pine can create leachates with very similar characteristics for oxygen level, pH, and conductivity [38]. In contrast, the biotic index for macroinvertebrates (IBMWP), was significantly higher in ponds in native forests than in eucalypt plantations. This index is designed to assess water quality [39] which suggests that eucalypt plantations have more toxic pond water. This conclusion is in line with the much higher concentration of phenolic compounds in eucalypt leachates than in those from other species such as alder or pine [38]. Finally, water conductivity also predicted amphibian richness which suggests that the accumulation of chemicals plays a role in amphibian diversity.

Anuran larvae are detritivore feeders, which makes them very sensitive to the quality of plant remains [40]. Detritus is known to be of lower quality in ponds in plantations than in native deciduous forest [21,41]. This could explain the reduced abundance of *R. temporaria* tadpoles in plantations compared to oak forests. If tadpole survival is lower in plantations due to toxicity of their food resources, the number of individuals that reach adulthood will be lower, reducing the population each generation. Moreover, poor environmental conditions and a low quality diet early in development can reduce adult size, lower energy reserves, decrease competitive ability and, ultimately, reduce fitness [42]. This could lower the ability of some adults to breed, and exacerbate the effect of a smaller adult population size.

### Plantation effects on habitat quality

Ponds in oak natural forests were significantly bigger and deeper than those in plantations, and the IBMWP is known to be lower in ponds that suffer periodic desiccation [43]. The periodic drying out of smaller ponds could further explain the observed differences in both the diversity and abundance of amphibians among the three habitat types. Interestingly, *R. dalmatina* and *T. marmoratus* have a long larval period of about 3 months that extends until summer [44]; and for *A. obstreticans* the tadpoles overwinter in the water, extending the larval cycle to a year [44]. These three species were absent in plantations, possibly because they cannot complete metamorphosis there before the ponds dry out. Both the accumulation of toxic compounds of leachates and the desiccation of the ponds in summer is expected to increase with on-going climate change.

The rapid drying out of smaller ponds might also explain the apparent habitat differences in aquatic vegetation: coverage is almost absent in ponds in pine plantations. The amount of aquatic vegetation in ponds might explain some of the variation in the density and abundance of amphibians. Amphibian species richness and individual density has been observed to increase with structural complexity of the habitat [45]. Terrestrial vegetation can provide refuge and feeding opportunities for anuran adults. Submerged vegetation provides refuge, food [46] and protection against UV-B radiation. In newts, aquatic vegetation is likely to be related to the oviposition behavior of wrapping each egg individually in leaves to protect them from UV-B radiation and predators [47].

### Conclusions

In sum, the land use seems to be a critical factor behind the differences we found in amphibian diversity and density among ponds. However, some related variables, such as water quality, macroinvertebrate availability (an index of food resources), and the early desiccation of the ponds in tree plantations also play an important role explaining the differences found among habitats. These effects seemed to be more detrimental for some species (*A. obstetricans, R. temporaria, R. dalmatina* and *T. marmoratus*) than for others (*L. helveticus* and *P. perezi*). Discovering which are the key drivers modulating the populations of amphibians under different land use regimes would allow to preserve the natural biodiversity of this threatened animal group.

## Acknowledgements

We thank Michael Jennions for his invaluable help improving the manuscript, and Ion Garin-Barrio, June Garrido, Javier Urquijo, Adolfo Iglesias, Oier Virizuela and Flora Carrasco for field assistance.

### Fundings

This work was supported by Basque Government and the Spanish Ministry of Education and Culture with a pre-doctoral grant to M. I-C (grant number FPU12/04148).

Amphibian data were collected following all Spanish legal requirements and the study was performed under license from the Álava, Bizkaia and Gipuzkoa Administrations. All capture and handling of individuals complied with the contemporary laws regulating the treatment of animals in Spain. MIC conducted this work with Spanish accreditation to conduct experiments with animals R.D 53/2013 (ref number 10/096442.9/13).

## References

1. MacDicken K. Global Forest Resources Assessment 2015: What, why and how? For Ecol Manage. 2015;352: 3–8. doi: 10.1016/j.foreco.2015.02.006

2. Buongiorno J, Zhu S, Zhang D, Turner J, Tomberlin D. Overview of the Global Forest Products. In: Elsevier Science, editor. The Global Forest Products Model (GFPM): Structure, Estimation, Applications. USA; 2003. pp. 252–262.

3. Lindenmayer D, Hobbs R. Fauna conservation in Australian plantation forests–a review. Biol Conserv. 2004;119: 151–168. doi: 10.1016/j.biocon.2003.10.028

4. Fork S, Woolfolk A, Akhavan A, Van Dyke E, Murphy S, Candiloro B, et al. Biodiversity effects and rates of spread of nonnative eucalypt woodlands in central California. Ecol Appl. 2015;25: 2306–2319. doi:10.1890/14-1943.1

5. Zurita G, Rey N, Varela D, Villagra M. Conversion of the Atlantic Forest into native and exotic tree plantations: Effects on bird communities from the local and regional perspectives. For Ecol Manage. 2006;235: 164–173. doi:10.1016/j.foreco.2006.08.009

6. Brockerhoff E, Jactel H, Parrotta J, Quine C. Plantation forests and biodiversity: oxymoron or opportunity? Biodivers Conserv. 2008;17: 925–951. doi:10.1007/978-90-481-2807-5_1

7. Heer K, Helbig-Bonitz M. Effects of land use on bat diversity in a complex plantation–forest landscape in northeastern Brazil. J Mammal. 2015;96: 720–731. doi:http: 10.1093/jmammal/gyv068

8. Mortelliti A, Lindenmayer D. Effects of landscape transformation on bird colonization and extinction patterns in a large-scale, long-term natural experiment. Conserv Biol. 2015;29: 1314–1326. doi: 10.1111/cobi.12523

9. Galván I, Benayas J. Bird species in Mediterranean pine plantations exhibit different characteristics to those in natural reforested woodlands. Oecologia. 2011;166: 305–316. doi:10.1007/s00442-010-1849-0

10. Rieff G, Natal-da-Luz T, Sousa J. Collembolans and mites communities as a tool for assessing soil quality: effect of eucalyptus plantations on soil mesofauna biodiversity. Curr Sci. 2016;110: 713–719.

11. Oliveira J, Fernandes F, Ferreira M. Effects of forest management on physical habitats and fish assemblages in Iberian eucalypt streams. For Ecol Manage. 2016;363: 1–10. doi:10.1016/j.foreco.2015.12.011

12. Mortelliti A, Michael D, Lindenmayer D. Contrasting effects of pine plantations on two skinks: results from a large-scale “natural experiment”in Australia. Anim Conserv. 2015;18: 433–441. doi:10.1111/acv.12190

13. Tererai F, Gaertner M, Jacobs S. Eucalyptus invasions in riparian forests: effects on native vegetation community diversity, stand structure and composition. For Ecol Manage. 2013;297: 84–93. doi:10.1016/j.foreco.2013.02.016

14. Arntzen J. Drastic population size change in two populations of the golden-striped salamander over a forty-year period—Are Eucalypt plantations to blame? Diversity. 2015;7: 270–294. doi:10.3390/d7030270

15. Cruz J, Sarmento P, Carretero M, White P. Exotic fish in exotic plantations: A multi-scale approach to understand amphibian occurrence in the Mediterranean region. PLoS One. 2015;10.

16. Souto X, Gonzales L, Reigosa M. Comparative analysis of allelopathic effects produced by four forestry species during decomposition process in their soils in Galicia (NW Spain). J Chem Ecol. 1994;20: 3005–3015. doi:10.1007/BF02098405

17. Pozo J, Basaguren A, Elosegui A, Molinero J, Fabre E. Afforestation with *Eucalyptus globulus* and leaf litter decomposition in streams of northern Spain. Hydrobiologia. 1998;101: 373–374. doi:10.1023/A:1017038701380

18. Florence R. Cultural problems of Eucalyptus as exotics. Commonw For Rev. 1986;65: 141–163.

19. Ferreira V, Larrañaga A, Gulis V, Basaguren A. The effects of eucalypt plantations on plant litter decomposition and macroinvertebrate communities in Iberian streams. For Ecol Manage. 2015;355: 129–138. doi: 10.1016/j.foreco.2014.09.013

20. Ferreira V, Elosegi A, Gulis V, Pozo J. Eucalyptus plantations affect fungal communities associated with leaf-litter decomposition in Iberian streams. Arch für Hydrobiol. 2006;166: 467–490. doi:10.1127/0003-9136/2006/0166-0467

21. Martínez A, Larrañaga A, Miguélez A. Land use change affects macroinvertebrate community size spectrum in streams: the case of *Pinus radiata* plantations. Freshw Ecol. 2016;61: 69–79. doi:10.1111/fwb.12680

22. Canhoto C, Laranjeira C. Leachates of *Eucalyptus globulus* in intermittent streams affect water parameters and invertebrates. Int Rev Hydrobiol. 2007;92: 173–182. doi:10.1002/iroh.200510956

23. Houlahan J, Findlay C, Schmidt B, Meyer A. Quantitative evidence for global amphibian population declines. Nature. 2000;404: 752–755. doi:10.1038/35008052

24. Cushman S. Effects of habitat loss and fragmentation on amphibians: a review and prospectus. Biol Conserv. 2006;128: 231–240. doi:10.1016/j.biocon.2005.09.031

25. Rowe CL, Hopkins WA, Bridges C. Physiological ecology of amphibians in relation to susceptibility to natural and anthropogenic factors. In: Linder G, Krest S, Sparling D, editors. Amphibian Decline: An Integrated Analysis of Multiple Stressor Effects. Pensacola: SETAC press; 2003. pp. 9–57.

26. Watling J, Hickman C, Orrock J. Invasive shrub alters native forest amphibian communities. Biol Conserv. 2011;144: 2597–2601. doi: 10.1016/j.biocon.2011.07.005

27. Semlitsch R, Bodie J. Biological criteria for buffer zones around wetlands and riparian habitats for amphibians and reptiles. Conserv Biol. 2003;17: 1219–1228. doi:10.1046/j.1523-1739.2003.02177.x

28. Euskalmet. Climatología del País Vasco. 2011. http://www.euskalmet.euskadi.eus/s07-5853x/es/contenidos/informacion/car_latitud/es_7257/es_latitud.html

29. Tachet H, Richoux P, Bournaud M, Usseglio-Polatera P. Invertébrés d’eau douce : Systématique, biologie, écologie. CNRS, editor. Paris; 2010.

30. Alba-Tercedor J, Jáimez-Cuéllar P, Álvarez M, Avilés J, Bonada i Caparrós N, Casas J, et al. Caracterización del estado ecológico de ríos mediterráneos ibéricos mediante el índice IBMWP (antes BMWP’). Limnetica. 2002;21: 175–185. doi:http://hdl.handle.net/2445/32903

31. Alba-Tercedor J, Sánchez-Ortega A. Un método rápido y simple para evaluar la calidad biológica de las aguas corrientes basado en el de Hellawell (1978). Limnetica. 1988;4: 41–56.

32. Freda J. The influence of acidic pond water on amphibians: a review. Water Air Soil Pollut. 1986;30: 439–450. doi:10.1007/BF00305213

33. Oertli B, Joye D, Castella E, Juge R, Cambin D. Does size matter? The relationship between pond area and biodiversity. Biol Conserv. 2002;104: 59–70. doi: 10.1016/S0006-3207(01)00154-9

34. Orizaola G, Braña F. Oviposition behaviour and vulnerability of eggs to predation in four newt species (genus *Triturus*). Herpetol J. 2003;13: 121–124.

35. Brodman R, Ogger J, Bogard T, Long A. Multivariate analyses of the influences of water chemistry and habitat parameters on the abundances of pond-breeding amphibians. J Freshw Ecol. 2003;18: 425–436. doi:10.1080/02705060.2003.9663978

36. Raffel T, Rohr J, Kiesecker J. Negative effects of changing temperature on amphibian immunity under field conditions. Funct Ecol. 2006;20: 819–828. doi:10.1111/j.1365-2435.2006.01159.x

37. Lengendre P, Lengendre L. Numerical Ecology. 2nd ed. Elsevier Science, editor. Amsterdam; 1998.

38. Friberg N, Winterbourn M. Interactions between riparian leaves and algal/microbial activity in streams. Hydrobiologia. 1996;341: 51–56. doi:10.1007/BF00012302

39. Jáimez-Cuéllar P, Vivas S, Bonada N, Robles S. Protocolo GUADALMED (prece). Limnetica. 2002;21: 187–204.

40. Maerz J, Cohen J, Blossey B. Does detritus quality predict the effect of native and non‐native plants on the performance of larval amphibians? Freshw Biol. 2010;55: 1694–1704. doi:10.1111/j.1365-2427.2010.02404.x

41. Molinero J, Pozo J. Impact of a eucalyptus (*Eucalyptus globulus* Labill.) plantation on the nutrient content and dynamics of coarse particulate organic matter (CPOM) in a small stream. Hydrobiologia. 2004;528: 143–165. doi:10.1007/s10750-004-2338-4

42. Taborsky B. The influence of juvenile and adult environments on life-history trajectories. Proc R Soc B. 2006;273: 741–750. doi:10.1098/rspb.2005.3347

43. Attrill M, Rundle S, Thomas R. The influence of drought-induced low freshwater flow on an upper-estuarine macroinvertebrate community. Water Res. 1996;30: 261–268. doi:10.1016/0043-1354(95)00186-7

44. García-París M, Montori-Faura A, Herrero-Solans P. Amphibia: Lissamphibia. Consejo Superior de Investigaciones Científicas, editor. Fauna Ibérica. 2004. p. 640.

45. Vallan D. Effects of anthropogenic environmental changes on amphibian diversity in the rain forests of eastern Madagascar. J Trop Ecol. 2002;18: 725–742. doi:10.1017/S026646740200247X

46. Waringer-Löschenkohl A. An experimental study of microhabitat selection and microhabitat shifts in European tadpoles. Amphibia-Reptilia. 1988;9: 219–236. doi:10.1163/156853888X00314

47. Alarcos G, Ortiz M, Lizana M, Aragón A. La colonización de medios acuáticos por anfibios como herramienta para su conservación: el ejemplo de Arribes del Duero. Munibe. 2003;16: 114–127.

